# Daily and Intermittent Smoking Decrease Gray Matter Volume and Concentrations of Glutamate, Creatine, Myo-Inositol and *N*-acetylaspartate in the Prefrontal Cortex

**DOI:** 10.1101/2020.09.08.288050

**Authors:** Paul Faulkner, Susanna Lucini Paioni, Petya Kozhuharova, Natasza Orlov, David J. Lythgoe, Yusuf Daniju, Elenor Morgenroth, Holly Barker, Paul Allen

## Abstract

Cigarette smoking is still the largest contributor to disease and death worldwide. Successful cessation is hindered by decreases in prefrontal glutamate concentrations and gray matter volume due to daily smoking. Because non-daily, intermittent smoking also contributes greatly to disease and death, understanding whether infrequent tobacco use is associated with reductions in prefrontal glutamate concentrations and gray matter volume may aid public health. Eighty-five young participants (41 non-smokers, 24 intermittent smokers, 20 daily smokers, mean age ~23 years old), underwent ^1^H-magnetic resonance spectroscopy of the medial prefrontal cortex, as well as structural MRI to determine whole-brain gray matter volume. Compared to non-smokers, both daily and intermittent smokers exhibited lower concentrations of glutamate, creatine, *N*-acetylaspartate and myo-inositol in the medial prefrontal cortex, and lower gray matter volume in the right inferior frontal gyrus; these measures of prefrontal metabolites and structure did not differ between daily and intermittent smokers. Finally, medial prefrontal metabolite concentrations and right inferior frontal gray matter volume were positively correlated, but these relationships were not influenced by smoking status. This study provides the first evidence that both daily and intermittent smoking are associated with low concentrations of glutamate, creatine, *N*-acetylaspartate and myo-inositol, and low gray matter volume in the prefrontal cortex. Future tobacco cessation efforts should not ignore potential deleterious effects of intermittent smoking by considering only daily smokers. Finally, because low glutamate concentrations hinder cessation, treatments that can normalize tonic levels of prefrontal glutamate, such as N-acetylcysteine, may help intermittent and daily smokers to quit.

## Introduction

Tobacco smoking is one of the leading causes of disease and death worldwide, contributing to roughly 6 million deaths per year around the globe (1). While most tobacco-related negative health effects are attributed to the daily smoking of many cigarettes, a recent meta-analysis reported that smoking only one cigarette on most days carries a risk for cardiovascular diseases that is half to two-thirds of that from smoking 20 cigarettes every day (2). Importantly, while the number of daily smokers has decreased in recent years, the proportion of cigarette users who smoke intermittently (e.g. by limiting smoking to non-daily social contexts) may actually be increasing (3, 4, 5, 6). In fact, up to a third of all smokers in the U.S. smoke on a non-daily basis (7), yet many of these individuals struggle to quit (8). There is thus a need to develop therapies that reduce cigarette consumption in both daily and intermittent smokers.

To aid cessation efforts, some research groups have examined the glutamatergic mechanisms of tobacco use disorder (e.g. 9), and report that daily smoking is associated with glutamatergic alterations in the prefrontal cortex. For example, one group demonstrated that, compared to non-smokers, daily smokers exhibit a significantly lower density of the metabotropic glutamate receptor 5 in many cortical regions, including the prefrontal cortex (10). Using ^1^H-Magnetic Resonance Spectroscopy (^1^H-MRS), Durazzo and colleagues observed lower concentrations of glutamate, as well as lower concentrations of creatine, *N*-acetylaspartate and myo-inositol, in the right dorsolateral prefrontal cortex of 30 adult daily smokers compared to 35 adult non-smokers (11). Interestingly, another study reported no difference in prefrontal glutamate concentrations between 13 daily smokers, 9 former smokers and 16 non-smokers (12); however, it is likely that this lack of a group effect is partially due to the low sample size in this study. Unfortunately, the authors did not report results of group comparisons in terms of creatine, *N*-acetylaspartate or myo-inositol. While daily smoking may be associated with reductions in prefrontal glutamate (and indeed concentrations of these other metabolites), it is currently unknown whether there is a similar association in intermittent smokers. Because low concentrations of prefrontal glutamate are related to a greater likelihood of relapse during a quit attempt in daily smokers (13), understanding whether non-daily, infrequent smokers also exhibit altered concentrations of glutamate and other metabolites in the prefrontal cortex could aid cessation efforts.

Structural neuroimaging studies have also indicated that daily cigarette smoking may damage the prefrontal cortex. For example, using voxel-based morphometry (VBM) (14), Brody and colleagues demonstrated that compared to non-smokers, daily smokers exhibit lower gray matter densities in the bilateral prefrontal cortex (including the orbitfrontal cortex and anterior cingulate cortex) and right cerebellum (15). These results have been replicated by a number of studies (e.g. 16), including by a study of 659 non-smokers and 315 daily smokers (17). However, to date, no study has examined whether this structural profile is seen in intermittent smokers, or whether such low prefrontal volume is related to changes in glutamate concentrations. Because nicotine administration releases glutamate in the prefrontal cortex (18), and because excessive glutamate release can cause neuronal damage (19, 20), it is possible that chronic smoking-related glutamate release in the prefrontal cortex is associated with the observed decreases in prefrontal gray matter in daily smokers. Understanding whether low prefrontal gray matter volume is related to prefrontal glutamate (or indeed other brain chemicals) in both intermittent and daily smokers may also aid cessation efforts.

It is important to consider young smokers in these efforts. Myelination and synaptic pruning in the frontal lobes continue into early adulthood (21). Therefore, determining the association between smoking behaviours and both prefrontal glutamate and volume during young adulthood may aid development of therapies that alleviate the negative effects of smoking on brain and cognitive development. Further, because young smokers a) typically smoke intermittently, b) find it easier to quit smoking than older smokers, and c) transition from intermittent smoking to daily smoking around 18-21 years of age (22), targeting younger smokers may result in higher sustained cessation rates than targeting older smokers (23).

We therefore aimed to compare prefrontal glutamate, brain volume and the relationship of the two, between daily smokers, non-daily intermittent smokers and non-smokers, using data collected from two studies of relatively young adults (mean age ~23 years old). It was hypothesized that, compared to non-smokers, both daily and intermittent smokers would exhibit lower glutamate concentrations and gray matter volume in the prefrontal cortex, and that daily smokers would exhibit even lower prefrontal glutamate and gray matter volume than intermittent smokers. On the basis of findings presented by Durazzo et al (11), we also aimed to determine the effects of both daily and intermittent smoking on prefrontal creatine, *N*-acetylaspartate and myo-inositol, and the relationship between concentrations of these metabolites and gray matter volume. Our hypotheses regarding these metabolites were secondary to, but mirrored, our hypothesis pertaining to prefrontal glutamate.

## Materials and Methods

We report an analysis of data collected from two separate studies of relatively young participants (mean age 23.41 years old); both studies had the specific aim of collecting health-related MRI data in young adults. Both study protocols used the same MRI sequences, and all data were acquired on the same 3T MRI scanner at the Combined Universities Brain Imaging Centre.

### Participants

Across both studies, eighty-five participants were recruited via print and online advertisements. Thirty-six of these subjects participated in study 1, and 49 subjects participated in study 2. All participants gave written informed consent after receiving a detailed explanation of study procedures (approved by the University of Roehampton Research Ethics committee). Exclusion criteria for both studies were: self-report of psychiatric comorbidity; current drug use/abuse or dependence (other than tobacco use disorder or cannabis use); history of neurological injury or disease, pregnancy and contraindications for MRI (e.g. metal implants). Twenty participants were defined as daily smokers because they self-reported smoking at least 5 cigarettes every day for at least one year, in line with definitions in previous research (e.g. 5, 6, 8, 25, 26, 27, 28); twenty-four participants were defined as intermittent smokers because they smoked between 1-4 cigarettes on at least 1 day per week; forty-one participants were defined as non-smokers because they smoked 0 cigarettes per day. Data from study 1 were collected from November 2015 – March 2018, while presented data from study 2 were collected from September 2017-August 2019.

### Questionnaire Measures

All participants completed a demographics form (developed in-house) to determine age, gender, level of education, daily tobacco use, number of months of smoking and daily cannabis use, as well as self-reported psychiatric comorbidity, neurological disorder or use of illicit drugs. Lifelong exposure to cigarettes was inferred from ‘pack years’, calculated as the average number of packs of cigarettes smoked per day multiplied by the number of years of smoking, as in previous research (e.g. 11, 12).

### ^1^H-MRS Data Acquisition, Pre-processing and Analysis

Of the 85 participants, two daily smokers did not undergo the ^1^H-MRS scan, meaning that metabolite concentrations were quantified from 83 participants (41 non-smokers, 24 intermittent smokers, 18 daily smokers). All ^1^H-MRS scans were acquired using the same 3T Siemens Magnetom TIM Trio MRI system using a 32-channel head coil. ^1^H-MRS *in vivo* spectra were acquired from the same 20 × 20 × 20mm voxel located in the right medial prefrontal cortex (typical location shown in Fig. 1A). The structure and function of the medial prefrontal cortex are related to the clinical features of tobacco use disorder (e.g. 29); this voxel placement therefore allowed us to test our own hypotheses pertaining to the effects of smoking on prefrontal metabolite concentrations. A medial position was also chosen as lateral voxels can be harder to place due to tissue boundaries. The voxel was placed manually by referring to the individual subject’s T1-weighted (MPRAGE) scan. Specifically, we ensured that we placed the voxel very close to the mid-line of the brain, and as anterior as possible whilst avoiding any gyri and cerebrospinal fluid. This meant that the voxel was placed both anterior and slightly dorsal to the corpus callosum, as can be seen in the representative placement shown in Fig. 1A. Spectra were acquired using a Spin ECho full Intensity-Acquired Localized spectroscopy (SPECIAL; 30) ^1^H-MRS sequence with water suppression (TR = 3000ms; TE = 8.5ms; Phase cycle Auto; 192 averages from the right prefrontal cortex voxel (31). Water un-supressed spectra (16 averages) were also acquired. Outer volume suppression slabs were applied 5mm from the edge of each side of the voxel (six slabs in total), both to suppress signals originating outside of the right medial prefrontal voxel, and to minimize motion artefact effects on spectra within the voxel.

**Figure 1.**
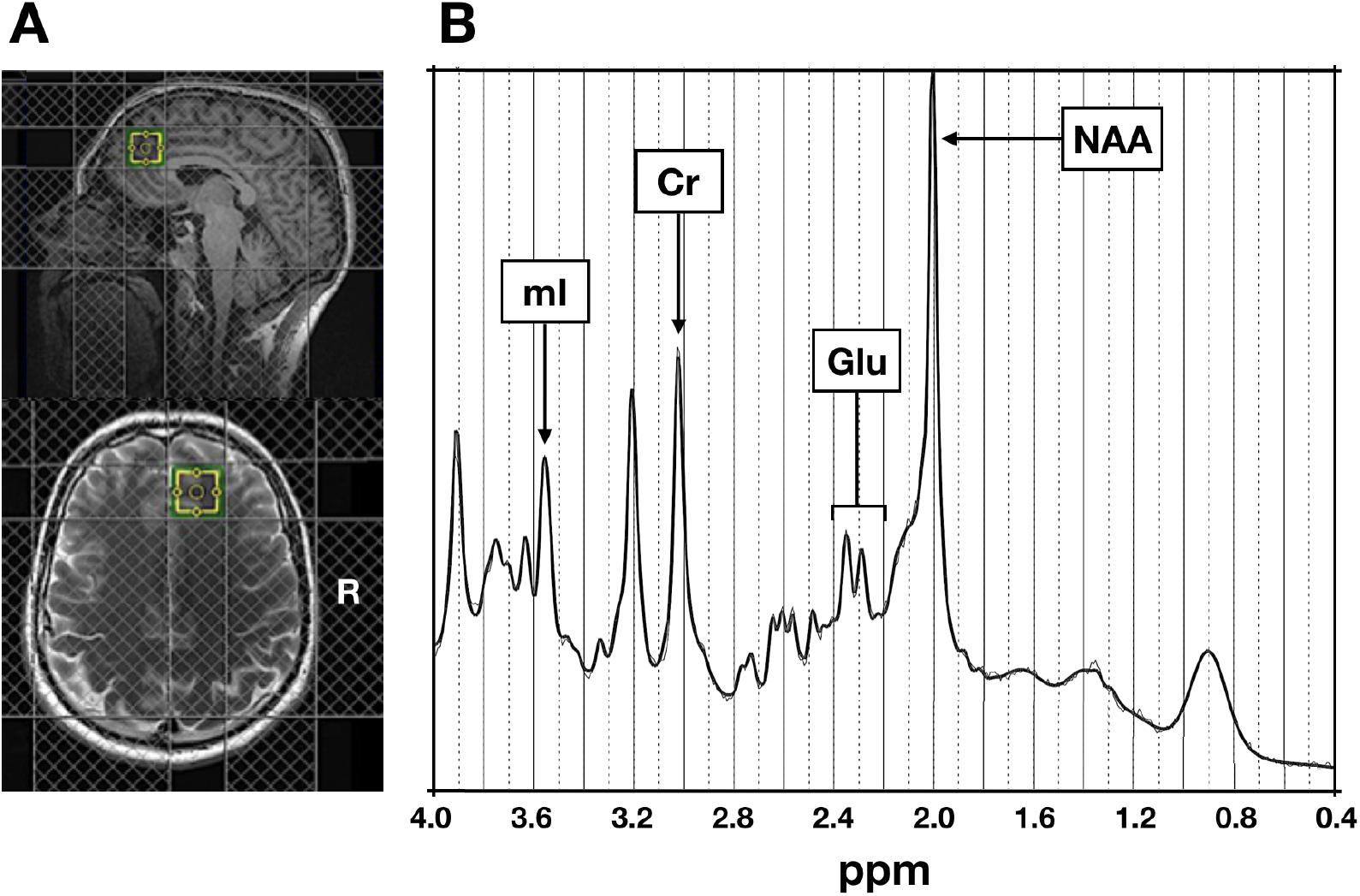
**A:** Typical ^1^H-MRS voxel placement in the medial prefrontal cortex. **B:** Example attained spectrum from the medial prefrontal voxel seen in A.

Spectra were analysed using LCModel 6.3-1L, with a basis set consisting of 19 simulated spectra; alanine (Ala), ascorbate (Asc), aspartate (Asp), creatine (Cr), γ-aminobutyric acid (GABA), glucose (Glc), glutamine (Gln), glutamate (Glu), glycine (Gly), glutathione (GSH), glycerophosphocholine (GPC), phosphocholine (PCh), lactate (Lac), myo-inositol (mI), N-acetylaspartate (NAA), N-acetylaspartateglutamate (NAAG), phosphorylethanolamine (PE), scyllo-inositol (Scyllo) and taurine (Tau). This basis set was simulated using FID-A (32) for TE = 8.5ms, magnetic field strength = ~3T and assuming ideal RF pulses. We excluded all spectra that had Cramér-Rao lower bounds of less than 20%. Line widths and signal-to-noise ratios were estimated as less than 8Hz and greater than 40, respectively (31). Cramér-Rao lower bounds (33), line widths and signal-to-noise ratios did not differ between daily smokers, intermittent smokers and non-smokers, or between study 1 and study 2 (see supplementary materials).

Water referencing and eddy current correction were used to quantify metabolite levels. When quantified in this way, such levels are influenced by cerebral spinal fluid, gray and white matter volumes of the region in which spectra are obtained (34), as well as by individual differences in whole-cortical gray matter (35). We therefore corrected these metabolite levels for gray and white matter content within the right medial prefrontal voxel using the GABA Analysis Toolkit (Gannet 3.1, http://gabamrs.blogspot.co.uk/), adapted to work with Siemens SPECIAL data. Segmentation was performed using ‘new segment’ in SPM12 (www.fil.ion.ac.uk/spm/software/spm8/). Cerebrospinal fluid, gray and white matter volumes were then accounted for in the expression of glutamate (Glu) using LCModel (36, 37). Specifically, Glu was corrected using the formula (Glu*(43300*gray matter volume + 35880*white matter volume + 55556*cerebrospinal fluid))/(35880*(1-cerebrospinal fluid)); glutamate concentrations corrected in this manner are denoted as ‘Glu *Corr*’. Concentrations of the remaining metabolites were also corrected using the same formula, and are referred to as the name of the metabolite followed by the suffix ‘*Corr*’, as in ‘Glu *Corr*’.

The effect of smoking status on concentrations of glutamate, creatine, myo-inositol and *N*-acetylaspartate in the prefrontal cortex was determined using separate ANOVAs, with the relevant metabolite added as the dependent variable and group added as a categorical factor; even though we did not have any hypotheses pertaining to the effect of smoking on concentrations of the remaining 15 metabolites quantified using ^1^H-MRS, we still examined the effect of smoking status on these metabolites for completeness (see supplementary materials). Because brain glutamate concentrations are influenced by gender (38), age (11), and daily cannabis use (39), these three variables were added to each model to control for their influence on each metabolite. To correct for the number of multiple comparisons, Fisher’s least significance difference method was used; specifically, once a main effect of smoking status on a metabolite was observed from the omnibus ANOVA that contained all three groups, pairwise comparisons (that controlled for the same variables as the omnibus test) were performed to determine whether each metabolite differed between a) non-smokers and daily smokers, b) non-smokers and intermittent smokers and c) intermittent smokers and daily smokers. Associations between each metabolite and indices of nicotine dependence (cigarettes per day and pack years) were examined using bivariate correlations.

### sMRI Data Acquisition, Pre-processing and Analysis

In both studies, high-resolution structural images were acquired using a T1-weighted magnetization-prepared rapid gradient echo (MPRAGE) sequence. Images were analysed using Computational Anatomy Toolbox 12 (CAT12; http://www.neuro.uni-jena.de/cat) implemented in SPM12 (Wellcome Trust Centre for Neuroimaging; www.fil.ion.ac.uk/spm/software/spm12). As per standard protocol (see http://www.neuro.uni-jena.de/cat12/CAT12-Manual.pdf), data were skull-stripped using the adaptive probability region-growing approach, normalized to the standard tissue probability map and segmented into gray matter, white matter and cerebral spinal fluid. These images were ‘modulated normalized’ images (i.e. voxel values were modulated using the Jacobian determinant), derived from the spatial normalization so that the absolute volume of gray matter could be compared between groups. This type of modulation requires group analyses to correct for individual differences in brain size; total intracranial volume was therefore added as a covariate to all group-level general linear models. The gray matter tissue segments then underwent statistical quality control testing for inter-subject homogeneity and overall image quality as included in the CAT12 toolbox, before manual visual inspection for potentially newly-introduced artefacts. These images were then registered to the MNI template using DARTEL registration and smoothed using an 8mm Gaussian Kernel.

Group level analyses were then performed by constructing one general linear model (GLM) in SPM12 that contained each participant’s modulated, normalized, segmented, registered and smoothed gray matter tissue segments. This GLM contained three explanatory variables (1 per group), along with separate variables for age, gender, daily cannabis use and total intracranial volume to control for the influence of these variables. Once a significant effect of group was identified from this F-test, whole-brain pairwise t-tests were constructed to determine brain regions in which gray matter was a) greater in non-smokers than in daily smokers; b) greater in non-smokers than in intermittent smokers; and c) greater in intermittent smokers than in daily smokers. While we hypothesised that smokers would exhibit *lower* prefrontal gray matter volume, we also constructed, for completeness, the opposite contrasts. A threshold of *p* < 0.05 with FWE correction for multiple comparisons was applied to all contrasts.

Finally, relationships between prefrontal glutamate and prefrontal volume were determined by using bivariate correlations. Specifically, these models included one of the metabolite *Corr* values, along with values from significant clusters identified from the above volumetric contrasts.

All ANOVAs and correlational analyses were performed using both frequentist and Bayesian analyses. Frequentist analyses were performed using the Statistical Package for Social Scientists version 26 (SPSS Inc., Chicago, Illinois). A significance threshold of alpha = 0.05 (2-tailed) was adopted. To correct for the number of comparisons due to the number of metabolites, a Bonferroni correction was performed. However, because our main hypotheses pertained to the effects of smoking status on prefrontal glutamate *only,* and because examinations of the remaining three metabolites were performed simply to corroborate the findings of (11) (and therefore were considered secondary and more exploratory), we only corrected for the three group comparisons of creatine, myo-inositol and *N*-acetylaspartate. We did not correct for the number of multiple tests due to testing the 15 remaining metabolites as these were only examined for completeness, and we had no reason to believe that smoking status would have an effect on these metabolites (see supplementary materials for these results). Bonferroni-corrected *p* values are denoted as ‘*P*_BC_’.

Bayesian analyses were performed using JASP (JASP Team (2019), version 0.11.1). Bayesian analyses were performed because they provide Bayes Factors, which depict a ratio of the probability of the evidence for one hypothesis (i.e. the null) relative to another (i.e. the experimental). This approach allows one to demonstrate support for the null hypothesis, as opposed to simply failing to reject the null as when using frequentist approaches (40). For these analyses, on the basis of (41), we considered Bayes Factors (*BF*_10_; NB: not logarithmically transformed) smaller than 1/100 to be extreme evidence for the null hypothesis, a *BF*_10_ between 1/100 and 1/30 to be very strong evidence for the null, a *BF*_10_ between 1/30 and 1/10 to be strong evidence for the null, a *BF*_10_ between 1/10 and 1/3 to be moderate evidence for the null, and a *BF*_10_ between 1/3 and 1 to be not worth more than a bare mention. Conversely, we considered *BF*_10_ larger than 100 to be extreme evidence for the experimental hypothesis, a *BF*_10_ between 100 and 30 to be very strong evidence for the experimental hypothesis, a *BF*_10_ between 30 and 10 to be strong evidence for the experimental hypothesis, a *BF*_10_ between 10 and 3 to be moderate evidence for the experimental hypothesis, and a *BF*_10_ between 3 and 1 to be not worth more than a bare mention.

## Results

### Participant Characteristics

Of the 85 participants across both studies, 46 were male and 39 were female (mean age = 23.41 years, SD = 4.46 years). Daily smokers smoked an average of 11.45 (SD = 4.73) cigarettes a day and demonstrated a mean pack years of 6.21 (SD = 5.38), while intermittent smokers smoked an average of 1.96 (SD = 1.21) cigarettes per day and demonstrated a mean pack years of 0.45 (SD = 0.32). Gender did not influence the number of cigarettes smoked per day or pack years in either smoking group (all *p*s > 0.428). There was no difference between groups in terms of age or gender (both *p*s > 0.109, both *BFs* < 1.264), although there was a trend towards the three groups differing in daily cannabis use (*F*(2,84) = 3.058, *p* = 0.053, *BF* = 2.890), with daily smokers tending to use more cannabis than non-smokers. A full summary of participant characteristics can be observed in Table 1. A description of all participant characteristics, is given in the supplementary materials.

**Table 1.** Participant Characteristics. ^a^ Denotes mean (SD). ^b^ Denotes Comparison of Daily Smokers and Intermittent Smokers Only.^c^ Denotes Mean (SD) Joints Per Day.

|  | Non-Smokers | Intermittent Smokers | Smokers | Group Comparison |
| --- | --- | --- | --- | --- |
| *N* | 41 | 24 | 20 | - |
| Gender (*M/F*) | 26/15 | 8/16 | 12/8 | $p = 0.109$ |
| Age (*years*)<sup>*a*</sup> | 22.87 (4.60) | 22.67 (3.75) | 25.40 (4.58) | $p = 0.122$ |
| Education (*years*)<sup>*a*</sup> | 15.14 (5.07) | 16.25 (1.59) | 16.69 (2.12) | $p = 0.315$ |
| Cigarettes Per Day<sup>*a*</sup> | - | 1.96 (1.21) | 11.45 (4.73) | $p < 0.001$<sup>*b*</sup> |
| Pack Years<sup>*a*</sup> | - | 0.45 (0.32) | 6.21 (5.37) | $p < 0.001$<sup>*b*</sup> |
| Cannabis Use<sup>*c*</sup> | 0.65 (0.84) | 1.03 (1.04) | 1.16 (1.16) | $p = 0.053$ |

### Influence of Smoking Status on Prefrontal Glutamate and Other Metabolites

There was a main effect of smoking status on Glu *Corr* values (*F*(2,77) = 6.188, *p* = 0.003, *BF* = 5.591; see Fig. 2A). Both daily and intermittent smokers exhibited significantly lower Glu *Corr* values in this region than non-smokers (daily smokers vs. non-smokers: *F*(1,54) = 6.530, *p* = 0.013; *BF* = 35.361; intermittent smokers vs. nonsmokers: *F*(1,60) = 6.109, *p* = 0.016, *BF* = 38.829). However, these values did not differ between daily smokers and intermittent smokers (*F*(1,37) = 0.057, *p* = 0.812, *BF* = 0.010).

**Figure 2.**
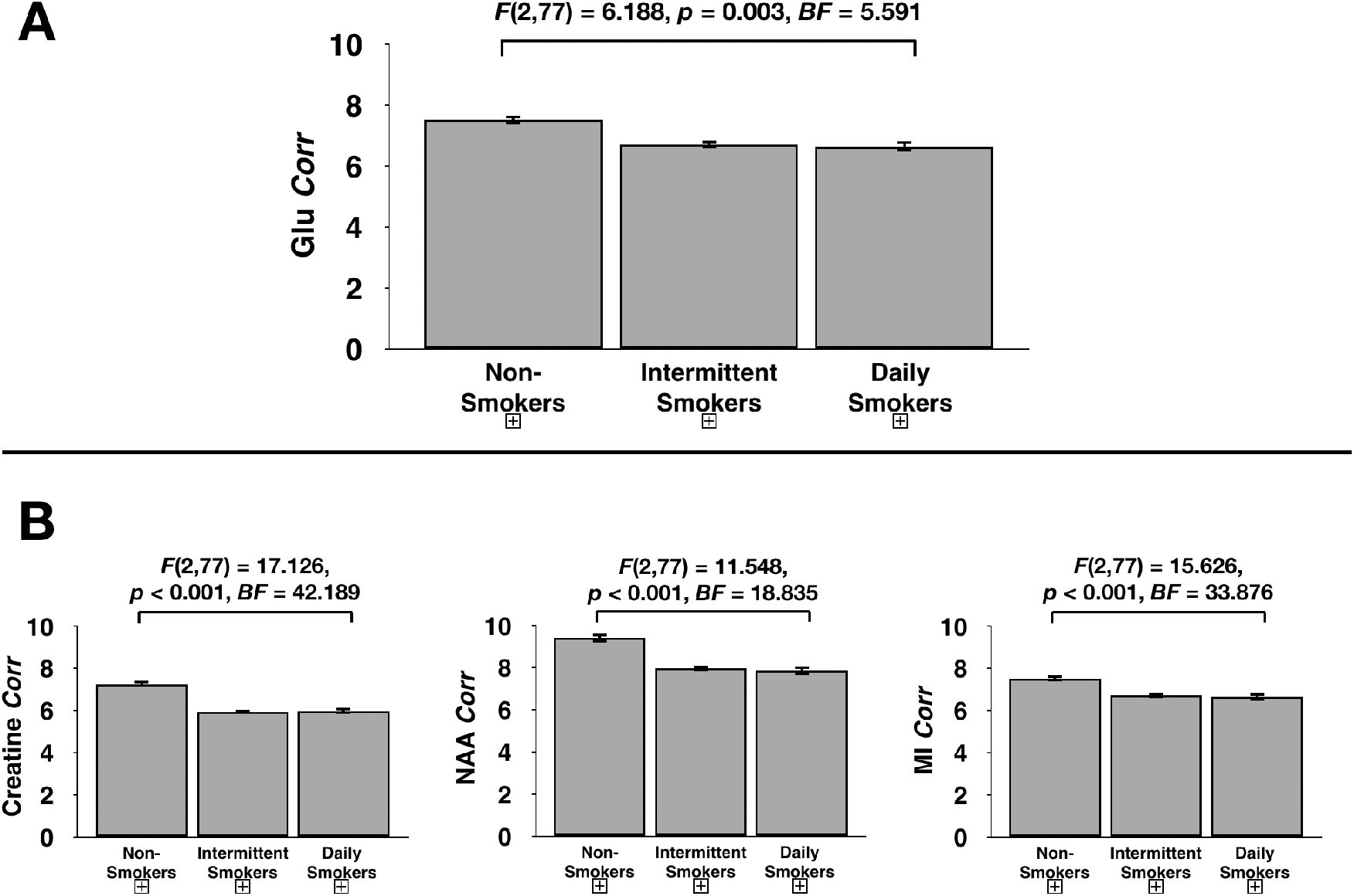
Glutamate **(A)** and creatine, N-acetylaspartate and myo-inositol **(B)** corrected for gray matter, white matter and cerebrospinal fluid volume within the voxel depicted in Figure 1A for non-smokers, intermittent smokers and daily smokers. Statistics are taken from the omnibus frequentist and Bayesian ANOVAs, and thus denote a meaningful difference between the three groups. Subsequent analyses revealed that both daily and intermittent smokers exhibited lower metabolite concentrations than non-smokers, but that daily and intermittent smokers did not differ in this variable.

There was also a main effect of smoking status on Creatine *Corr* values (*F*(2,77) = 17.126, *p* < 0.001, *P*_BC_ < 0.001, *BF* = 42.189), on NAA *Corr* values (*F*(2,77) = 11.548, *p* < 0.001, *P*_BC_ < 0.001, *BF* = 18.835) and on MI *Corr* values (*F*(2,77) = 15.626, *p* < 0.001, *P*_BC_ < 0.001, *BF* = 33.876) (see Fig. 2B). Subsequent analyses revealed that compared to non-smokers, both daily and intermittent smokers exhibited lower concentrations of each of these three metabolites (all *P*_BC_ < 0.008, all *BF*s 12.537), but that these concentrations did not differ between daily and intermittent smokers (all *p*s > 0.507, all *BF*s < 0.328). There were no effects of smoking status on any of the remaining 15 metabolites; a full description of these results can be seen in the Supplementary Materials.

Bivariate correlation analyses revealed that in the whole group, there was a slight negative correlation between age and Glu *Corr* values (*r* = −0.263, *p* = 0.018, *BF* = 4.281, see Fig. S2A), but not with Creatine *Corr* values, MI *Corr* values or NAA *Corr* values (all *p*s > 0.279, all *BF*s < 0.486). An independent samples t-test revealed that gender did not influence Glu *Corr* values (*t*(81) = 1.458, *p* = 0.149, *BF* = 0.242, see Fig. S2B), Creatine *Corr* values (*t*(81) = 1.322, *p* = 0.187, *BF* = 0.247), NAA *Corr* values (*t*(81) = 1.521, *p* = 0.132, *BF* = 0.855) or MI *Corr* values (*t*(81) = 1.412, *p* = 0.162, *BF* = 0.820). Finally, there was no correlation between the amount of cannabis smoked per day and Glu *Corr* values (*r* = −0.020, *p* = 0.910, *BF* = 0.192; see Fig. S3), Creatine *Corr* values (*r* = 0.037, *p* = 0.829, *BF* = 0.378), MI *Corr* values (*r* = 0.139, *p* = 0.420, *BF* = 1.138) or NAA *Corr* values (*r* = −0.053, *p* = 0.757, *BF* = 0.191).

### Influence of Smoking Status on Brain Volume

The whole-brain *F*-test revealed a significant main effect of smoking status on gray matter volume in a cluster of 673 voxels within the right inferior frontal gyrus (Fig. 3A); the local maxima of this cluster was *x* = 48, *y* = 36, *z* = −10. The whole-brain pairwise *t*-tests subsequently revealed that compared to non-smokers, both daily smokers and intermittent smokers exhibited lower gray matter volume in a cluster of 798 and 126 voxels, respectively, within the right inferior frontal gyrus; the local maxima of these clusters for both contrasts was identical to that of the cluster from the whole-group *F*-test (*x* = 48, *y* = 36, *z* = −10) (Figs. 3B and 3C). The remaining pairwise comparisons revealed no suprathreshold clusters.

**Figure 3.**
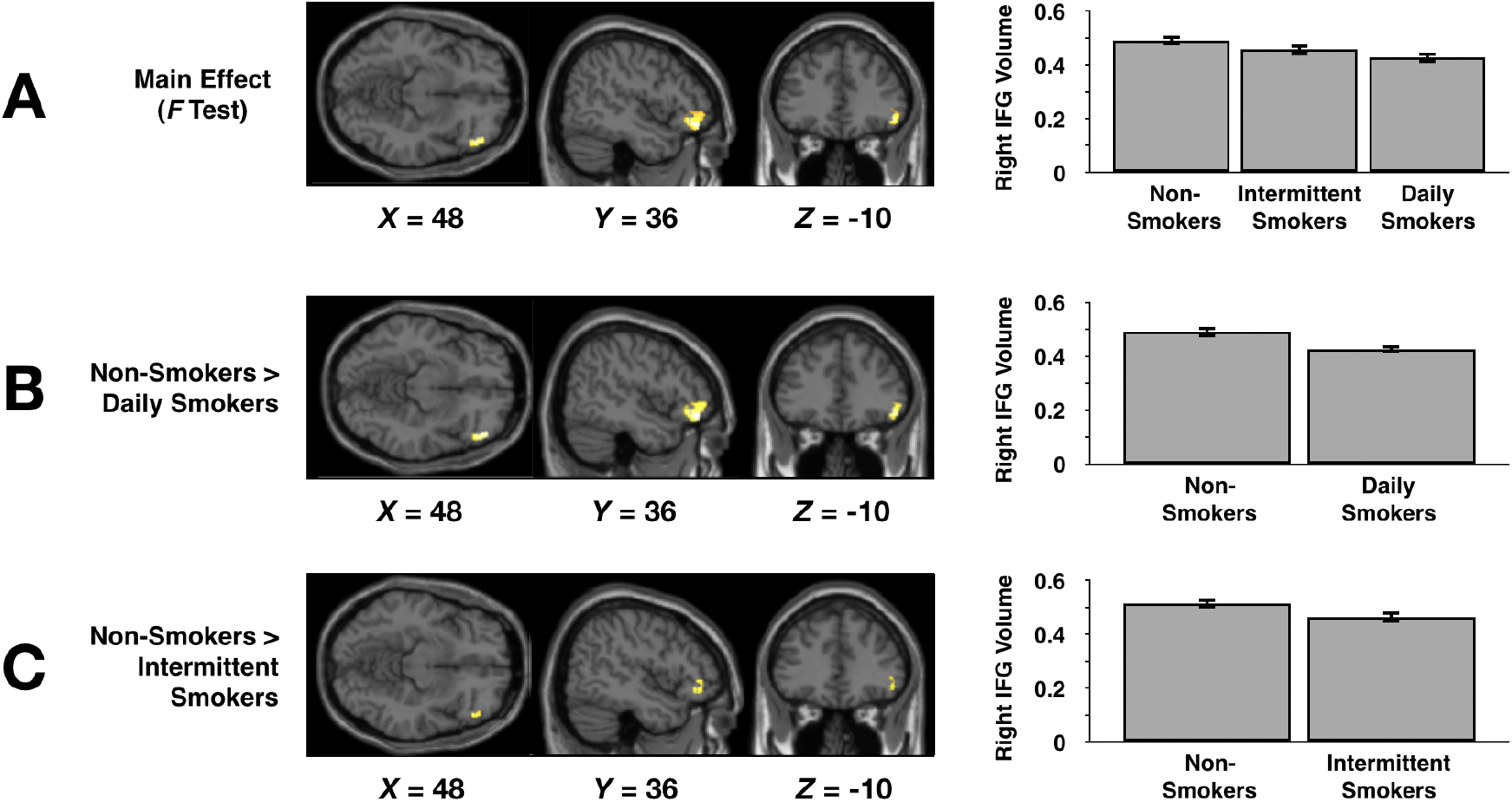
**A:** Results of the whole-group F-test that examined brain regions in which gray matter differed between the three groups. **B:** Results of the pair-wise t-test that examined brain regions in which gray matter differed between the daily smokers and non-smokers. **C:** Results of the pair-wise t-test that examined brain regions in which gray matter differed between the intermittent smokers and non-smokers. Bar graphs denote values extracted from the yellow cluster of voxels in the right inferior frontal gyrus in the relevant statistical parametric map on the left.

### Relationship of Brain Volume with Prefrontal Glutamate and Other Metabolites

A bivariate correlation revealed a significant positive association between Glu *Corr* values and values extracted from the cluster of 673 voxels depicted in Figure 3A (values averaged across all voxels in the cluster) (*r* = 0.244, *p* = 0.030, *BF* = 3.617) (see Fig. 4A). However, this relationship did not significantly differ between the three groups (*F*(2,77) = 0.158, *p* = 0.854, *BF* = 0.261).

**Figure 4.**
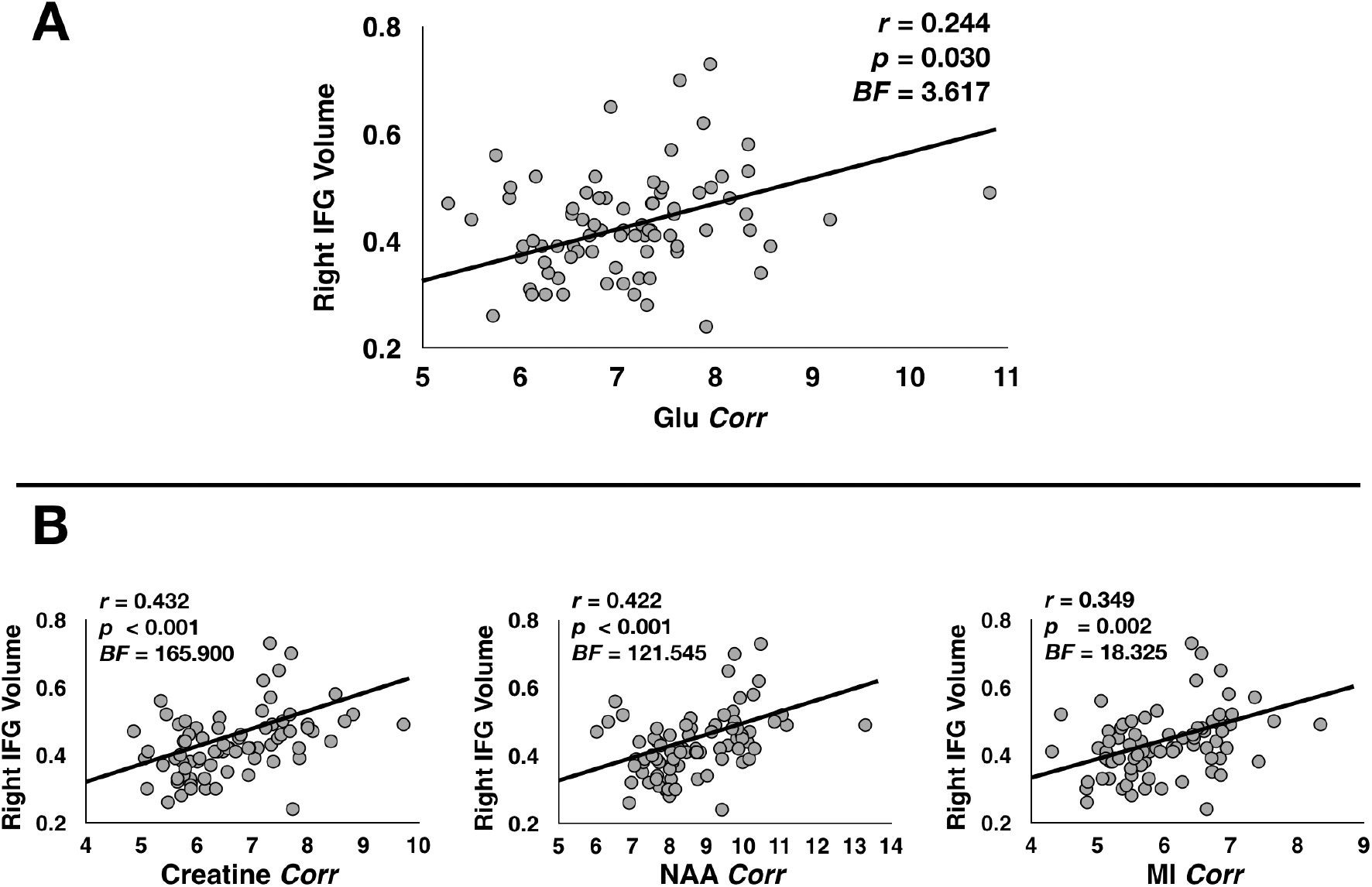
Correlation between metabolite concentrations in the voxel shown in Figure 1A, and gray matter volume in the cluster of voxels shown in Figure 3A in all participants. ‘Glu Corr’, ‘Creatine Corr’, ‘NAA Corr’ and ‘MI Corr’ denote gray matter volume-, white matter volume- and cerebrospinal fluid volume – corrected concentrations of glutamate **(A)** and creatine, N-acetylaspartate and myo-inositol **(B)**.

The values extracted from the cluster of 673 voxels depicted in Figure 3A also significantly and positively correlated with Creatine *Corr* values (*r* = 0.432, *p* < 0.001, *BF* = 165.900), NAA *Corr* values (*r* = 0.422, *p* < 0.001, *BF* = 121.545) and MI *Corr* values (*r* = 0.349, *p* = 0.002, *BF* = 18.325) (see Fig. 4B). None of these relationships were influenced by smoking status (all *p*s > 0.680, all *BF*s < 0.309).

## Discussion

This is the first study to examine brain metabolites and gray matter volume in non-daily, intermittent smokers. Our results suggest that, compared to non-smokers, both daily and intermittent smokers exhibit lower concentrations of glutamate, as well as lower concentrations of prefrontal creatine, myo-inositol and *N*-acetylaspartate, and lower gray matter volume in the prefrontal cortex. Interestingly, we found no difference between the two smoking groups in terms of these variables. Our findings extend those of previous research by indicating that even non-daily, intermittent smokers with relatively short smoking histories may exhibit low metabolite concentrations and gray matter volume in the prefrontal cortex, and that both daily and non-daily smoking over a shorter-than-average timespan may have therefore be related to abnormal prefrontal structure and chemistry.

Our primary hypothesis pertained to group differences in prefrontal glutamate and gray matter volume. The fact that both the daily and intermittent smokers exhibited significantly lower prefrontal glutamate and gray matter volumes than our aged-matched non-smokers was expected on the basis of findings that older adult smokers exhibit lower concentrations of glutamate in the dorsolateral prefrontal cortex compared to aged-matched non-smokers (11). However, the fact that there were no differences between the daily and intermittent smokers in these variables was contrary to our prediction that the former would exhibit lower glutamate and gray matter volume than the latter. Importantly, the relatively small sample size of 20 daily smokers and 24 intermittent smokers may have contributed to low power to detect a difference between these two groups in terms of prefrontal glutamate (and indeed creatine, *N*-acetylaspartate and myo-inositol). However, our Bayesian analyses provided ‘very strong’ evidence for the null hypothesis that prefrontal glutamate concentrations do not differ between these two groups at all, by providing a Bayes Factor of between 1/100 and 1/30 (BF = 0.010). Because our daily smokers smoked on average more than 11 cigarettes every day, these results might suggest that smoking on average 2 cigarettes per day on a non-daily, intermittent basis may have similar deleterious effects on prefrontal glutamate as smoking over half a pack of cigarettes every single day. Future studies could truly determine whether the similarity between these two groups is meaningful by comparing prefrontal metabolite concentrations in a large sample of daily and intermittent smokers.

The fact that smoking is associated with low glutamate and gray matter volume in the prefrontal cortex may be partially explained by the fact that nicotine acutely induces glutamate release in the prefrontal cortex (18), and that chronic elevated glutamate (due to repetitive smoking, for example) can promote cell death via excitotoxicity in this brain region (e.g. 19). Importantly, while excitotoxicity-induced cell death occurs due to *elevated* extracellular glutamate, the current findings suggest that smoking is associated with *decreases* in prefrontal glutamate concentrations, as do those results published previously (11). However, it is important to note that the excitotoxic effects of extracellular glutamate cannot be observed using ^1^H-MRS; this technique does not distinguish between intracellular and extracellular concentrations of glutamate, nor does it allow for observation of glutamate function at the level of the receptor. To truly determine the causal effects of smoking on brain glutamate throughout the lifespan, future studies could utilize other neuroimaging techniques such as Positron Emission Tomography, alongside ^1^H-MRS and structural MRI, to examine prefrontal glutamate concentrations, receptor density and gray matter volume at many stages throughout a smoker’s lifespan (e.g. 22).

These findings may have implications for the use of glutamatergic pharmacotherapies for smoking cessation. Research has shown that daily smokers who exhibit low glutamate concentrations in the dorsal anterior cingulate cortex prior to a quit attempt were more likely to relapse than those smokers who displayed higher glutamate concentrations in this region (13). This indicates that therapies which increase prefrontal glutamate concentrations may render smokers more able to quit smoking. One potential such therapy is N-acectylcysteine, which activates the cystine – glutamate exchanger to exchange extracellular cystine for intracellular glutamate and thus normalize levels of extracellular glutamate. Systemic administration of this drug has been shown to restore extracellular levels of glutamate and prevent drug-seeking behavior in rodents pre-treated with cocaine (42, 43). Further, two pilot studies with small sample sizes have shown that daily cigarette consumption decreases after administration of N-acetylcysteine for 4 weeks (44) and for 12 weeks (45). Future studies could therefore examine whether this drug, or indeed other promising glutamatergic pharmacotherapies such as Memantine or Topiramate (see 46), can increase prefrontal glutamate concentrations to aid smoking cessation over a longer period of time in a large sample of smokers.

Interestingly, results also largely supported our secondary hypothesis that smokers would exhibit lower concentrations of creatine, myo-inositol and *N*-acetylaspartate than non-smokers. However, contrary to our predictions, we found no differences between the daily and intermittent smokers in terms of these metabolites. Again, this may be due to the relatively small sample of daily and intermittent smokers, but our Bayesian analyses provided either ‘moderate’, ‘strong’ or ‘very strong’ evidence for the null hypothesis these two groups did not differ in terms of these metabolites in the prefrontal cortex. To better determine whether these two groups do not differ in terms of creatine, *N*-acetylaspartate and myoinositol, future studies could compare these metabolites in a much larger sample of daily and intermittent smokers. It is noteworthy that statistically speaking, group differences in concentrations of these three brain metabolites appear to be much larger than those in prefrontal glutamate. Indeed, studies have largely ignored the effects of cigarette smoking, daily or otherwise, on creatine, myo-inositol and *N*-acetylaspartate, yet studies that have examined such effects generally note that daily smokers exhibit lower concentrations of all three metabolites (e.g. 11, 12, 47, 48). While the functional and clinical relevance of such tobacco-related alterations are currently unclear, some research groups report that pharmacotherapies that act to increase levels of these metabolites may aid in drug cessation (e.g. 49). As such, one potential avenue for future research may be to examine the utility of pharmacotherapies that act on these brain metabolites. Finally, it is important to note that no effect of smoking status was observed on any of the remaining 15 metabolites that were quantified (e.g. such as GABA or glycine). This indicates that smoking may not influence all brain metabolites universally, but that cigarette use, whether on a daily or intermittent basis, may specifically affect concentrations of glutamate, creatine, *N*-acetylaspartate and myo-inositol. However, although these results support those of (11), future studies may wish to examine this hypothesis in a much larger sample of smokers.

The results of our VBM analyses replicate findings from previous studies (e.g. 15, 16, 17), by indicating that compared to non-smokers, smokers exhibit lower grey matter volume in the right inferior frontal gyrus. While our data do not indicate the functional relevance of smoking-related reductions in IFG gray matter volume, such reductions may be associated with known smoking-related deficits in cognitive control. For example, Tabibnia and colleagues examined brain activation during performance of motor, affective and craving self-regulation tasks in 25 daily smokers, and report that the IFG was commonly activated in all three tasks (50). Importantly, deficits in cognitive-control of emotional states are known to hinder smoking cessation (e.g. 51), and Faulkner et al recently reported that such deficits in smokers are related to weak connectivity of the IFG with the amygdala (52). As such, future studies could examine whether structural and functional abnormalities of the IFG contribute to difficulties in quitting smoking.

Although we report a significant positive correlation between prefrontal glutamate and gray matter volume, it is important to note that glutamate concentrations were quantified from the medial prefrontal cortex (see Fig. 1A), while gray matter volume was extracted from the region in which there was an effect of smoking status, the right inferior frontal gyrus. There was no justification to examine the relationship between glutamate and other metabolites with gray matter volume in the medial prefrontal region presented in Figure 1A, as there was no significant effect of smoking status on gray matter volume in this region. However, there is no reason to believe that gray matter volume in the inferior frontal gyrus should be uniquely and solely influenced by glutamate in the medial prefrontal cortex, and it is our belief that such glutamate concentrations may be associated with gray matter volume in many regions within the frontal cortex. In addition, our results indicate that the relationship between these two variables are not influenced by smoking behaviours. We believe that in light of the (expected) findings that smoking is associated with reductions in both glutamate concentrations and gray matter volume, that this lack of an effect makes statistical sense; if cigarette smoking had deleterious effects on glutamate and other metabolites but not gray matter volume (or vice versa), then this relationship would be influenced by smoking status. However, we again cannot rule out the notion that we lacked statistical power to detect such an effect; future studies could therefore examine this influence of smoking behaviours on the relationship between prefrontal glutamate and gray matter volume in a larger sample of participants.

This study has several limitations. First, because young smokers typically have lower levels of nicotine dependence, our results from young participants may not be generalizable to the wider population of cigarette users. Secondly, our relatively small sample size of 20 daily smokers limited our ability to determine the effects of other individual differences, such as those in gender and cigarette smoking behaviours. For instance, we observed a small-to-moderate strength negative correlation between prefrontal glutamate concentrations and both the average number of cigarettes smoked per day and pack years in daily smokers (r = ~−0.3 in both instances, which replicates the results of (11); see supplementary materials); however, both frequentist and Bayesian analyses revealed that the strength of these relationships were non-significant and not more worth more than a bare mention, likely due to the small number of daily smokers. Thirdly, we did not quantify the duration of smoking abstinence prior to testing, meaning that we cannot determine the influence of abstinence on study variables. Fourthly, our cross-sectional design did not allow for us to make strong predictions regarding any causal effects of cigarette smoking on brain metabolites. As such, future studies may wish to use longitudinal designs, allowing stronger causal inferences, to examine the effects of tobacco smoking on brain volume and chemistry. Further, a number of participants self-reported light-to-moderate use of cannabis. Although our analyses indicated that such drug use did not influence prefrontal metabolites or gray matter volume, there is a possibility that it had a small, yet undetectable, effect on these variables. Finally, tobacco-related alterations in prefrontal structure and function are considered to be related to smoking-induced cognitive deficits (e.g. 53). However, due to differences in protocols between the two studies reported, examinations of group differences in cognitive performance were not possible. Understanding the glutamatergic mechanisms of smoking-related cognitive deficits could further aid smoking cessation efforts however.

In summary, this study provides the first piece of evidence that non-daily, infrequent smokers exhibit abnormally low prefrontal glutamate, creatine, myo-inositol and *N*-acetylaspartate concentrations and gray matter volume, and that these low concentrations are very similar in magnitude to those observed in daily smokers. Future research is needed to achieve a more complete understanding of the effects of these prefrontal changes on smoking behaviours and cessation, as well as on brain function and cognition, particularly throughout the lifespan.

## Supporting information

Supplementary Results

## Acknowledgments

We wish to acknowledge Ali Lingeswaran at the Combined Universities Brain Imaging Centre (CUBIC) at the University of London, Royal Holloway, for MRI scanning and technical support.

## Authors Contribution

PF was chiefly responsible for data analysis and interpretation of findings, as well as the writing of the manuscript. SLP was responsible for data analysis, particularly of structural MRI data. PK provided intellectual input regarding analysis of structural MRI data and interpretation of findings. NO provided intellectual input regarding analysis of the spectroscopy data. DJL was chiefly responsible for analysing the spectroscopy data, and interpretation of findings. NO, YD, EM and HB were responsible for data acquisition. PA was responsible for the conception and design of the work, and gave significant input regarding the interpretation of findings and writing of the manuscript. All authors gave final approval of the manuscript to be published.

