## Supplementary Results for "Daily and Intermittent Smoking Decrease Gray Matter Volume and Concentrations of Glutamate, Creatine, Myo-Inositol and *N*-acetylaspartate in the Prefrontal Cortex"

**Supplementary Materials**

### ***Results***

#### *Participant Characteristics*

Of the 85 participants across both studies, 46 were male and 39 were female (mean age = 23.41 years, SD = 4.46 years). The three groups did not differ in terms of gender composition ($F(2,84) = 2.068$, $p = 0.109$, $BF = 0.947$) or age ($F(2,84) = 2.129$, $p = 0.122$, $BF = 1.263$). However, there was a trend towards smoking status influencing magnitude of daily cannabis use (mean daily cannabis use for; non-smokers = 0.55 (SD = 0.84) joints per day; intermittent smokers = 1.03 (SD = 1.04) joints per day; daily smokers = 1.14 (SD = 1.17) joints per day; $F(2,84) = 3.058$, $p = 0.053$, $BF = 2.891$).

Of 36 participants in study 1, 34 were defined as non-smokers, 1 as an intermittent smoker, and 1 as a daily smoker. Of the 49 participants in study 2, seven were defined as non-smokers, 23 as intermittent smokers, and 19 as daily smokers. Participants from study 1 and study 2 did not differ in terms of gender composition ($F(1,84) = 2.298$, $p = 0.134$, $BF = 1.198$), or mean age ($F(1,84) = 2.173$, $p = 0.144$, $BF = 1.124$). However, these participants differed in daily cannabis use (mean daily cannabis use for participants in; study 1 = 0.31 (SD = 0.64) joints per day; study 2 = 1.15 (SD = 1.01) joints per day; $F(1,84) = 4.378$, $p = 0.021$, $BF = 4.890$). Because of these results, cannabis use was added to all analyses, to control for the influence of this variable on the relevant dependent variable. For a summary of these data, please see table S1.

**Table1. Participant Characteristics**

| | Study 1 | Study 2 | *Group Comparison* |
|---|---|---|---|
| *N* [a] | 34/1/1 | 7/23/19 | - |
| Gender (*M/F*) | 25/11 | 21/28 | *p* = 0.134 |
| Age (*years*) [b] | 22.58 (4.74) | 24.04 (4.25) | *p* = 0.144 |
| Education (*years*) [b] | 14.94 (5.31) | 16.45 (1.82) | *p* = 0.105 |
| Cannabis Use [c] | 0.31 (0.64) | 1.15 (1.01) | *p* = 0.021 |

**Table 1.** *Participant Characteristics for study 1 and study 2.* [a] *Denotes number of non-smokers/intermitter smokers/daily smokers.* [b] *Denotes mean (SD).* [c] *Denotes Mean (SD) Joints Per Day.* [d] *Denotes Mean (SD) Drinks Per Week*

Daily smokers smoked an average of 11.45 (SD = 4.73) cigarettes a day and demonstrated a mean pack years of 6.21 (SD = 5.37), while intermittent smokers smoked an average of 1.96 (SD = 1.21) cigarettes per day and demonstrated a mean pack years of 0.45 (SD = 0.32). In smokers, there was no influence of sex/gender on cigarettes smoked per day (males: mean = 12.00, SD = 4.90; females: mean = 11.80, SD = 5.63; $F(1,19) = 0.011$, $p = 0.918$, $BF = 0.189$) or pack years (males: mean = 7.30, SD = 6.17; females: mean = 6.36, SD = 5.31; $F(1,19) = 0.019$, $p = 0.891$, $BF = 0.214$). Further, in intermittent smokers, there was no influence of sex/gender on cigarettes smoked per day (males: mean = 1.65, SD = 0.76; females: mean = 2.09, SD = 1.36; $F(1,21) = 0.651$, $p = 0.429$, $BF = 0.763$) or pack years (males: mean = 0.33, SD = 0.20; females: mean = 0.49, SD = 0.35; $F(1,21) = 0.302$, $p = 0.588$, $BF = 0.522$). Finally, the three groups did not significantly differ in levels of education ($F(2,74) = 1.152$, $p = 0.322$, $BF = 0.891$).

### *Influence of Smoking Status on Brain Metabolites*

In smokers, there was no association between Glu *Corr* and either average number of cigarettes smoked per day ($r = -0.332$, $p = 0.226$, $BF = 1.017$) or pack years ($r = -0.350$, $p = 0.201$ $BF = 1.246$). Further, in intermittent smokers, Glu *Corr* was not significantly correlated with either cigarettes per day ($r = 0.149$, $p = 0.353$, $BF = 0.282$) or pack years ($r = 0.149$, $p = 0.352$, $BF = 0.285$). These relationships can be seen in Figure S1.

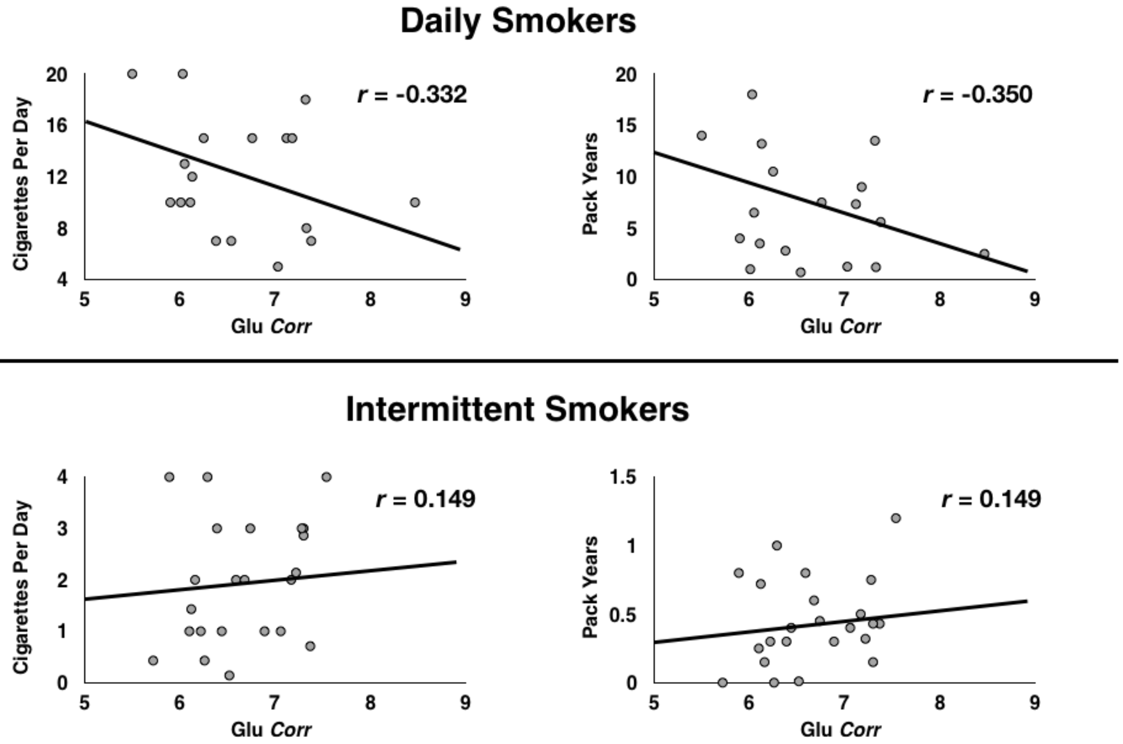


**Figure S1.** *Non-significant relationships between both cigarettes per day (left) and pack years (right) and glutamate concentrations in daily smokers (top) and intermittent smokers (bottom). 'Glu Corr' denotes glutamate concentrations corrected for gray matter, white matter and cerebrospinal fluid volume.*

The bivariate correlation analyses revealed that in smokers, there was no significant association between Glu *Corr* and either average number of cigarettes smoked per day ($r$ = -0.332, $p$ = 0.226, $BF$ = 1.017) or pack years ($r$ = -0.350, $p$ = 0.201, $BF$ = 1.246), between Creatine *Corr* and either average number of cigarettes smoked per day ($r$ = 0.081, $p$ = 0.779, $BF$ =0.143) or pack years ($r$ = 0.056, $p$ = 0.832, $BF$ = 0.106), between MI *Corr* and either average number of cigarettes smoked per day ($r$ = -0.137, $p$ = 0.599, $BF$ = 0.201) or pack years ($r$ = -0.194, $p$ = 0.488, $BF$ = 0.214), or between NAA *Corr* and either average number of cigarettes smoked per day ($r$ = -0.249, $p$ = 0.336, $BF$ = 0.443) or pack years ($r$ = -0.259, $p$ = 0.351, $BF$ = 0.459).

Further, in intermittent smokers, there was no significant association between Glu *Corr* and either average number of cigarettes smoked per day ($r$ = 0.149, $p$ = 0.353, $BF$ = 0.282) or pack years ($r$ = 0.149, $p$ = 0.352, $BF$ = 0.285), between Creatine *Corr* and either average number of cigarettes smoked per day ($r$ = 0.028, $p$ = 0.901, $BF$ = 0.096) or pack years ($r$ = 0.137, $p$ = 0.532, $BF$ = 0.207), between MI *Corr* and either average number of cigarettes smoked per day ($r$ = -0.001, $p$ = 0.998, $BF$ = 0.023) or pack years ($r$ = 0.177, $p$ = 0.420, $BF$ = 0.373), or between NAA *Corr* and either average number of cigarettes smoked per day ($r$ = 0.126, $p$ = 0.568, $BF$ = 0.229) or pack years ($r$ = 0.337, $p$ = 0.116, $BF$ = 1.442).

As expected, Glu *Corr* values negatively correlated with age in the whole group ($r$ = -0.265, $p$ = 0.017, $BF$ = 4.281) (Figure S2A), although this relationship did not reach statistical significance in each group alone (smokers: $r$ = -0.479, $p$ = 0.061, $BF$ = 2.772; intermittent smokers: $r$ = 0.142, $p$ = 0.507, $BF$ = 0.599; non-smokers: $r$ = -0.301, $p$ = 0.070, $BF$ = 2.268).

Gender did not influence Glu *Corr* values in the whole group ($F$(1,79) = 1.994, $p$ = 0.162, $BF$ = 0.950) (Figure S3B), or separately in smokers ($F$(1,19) = 1.062, $p$ = 0.320, $BF$ = 0.628), in chippers ($F$(1,22) = 0.053, $p$ = 0.821, $BF$ = 0.201) or in non-smokers ($F$(1,39) = 2.357, $p$= 0.133, $BF$ = 0.791).

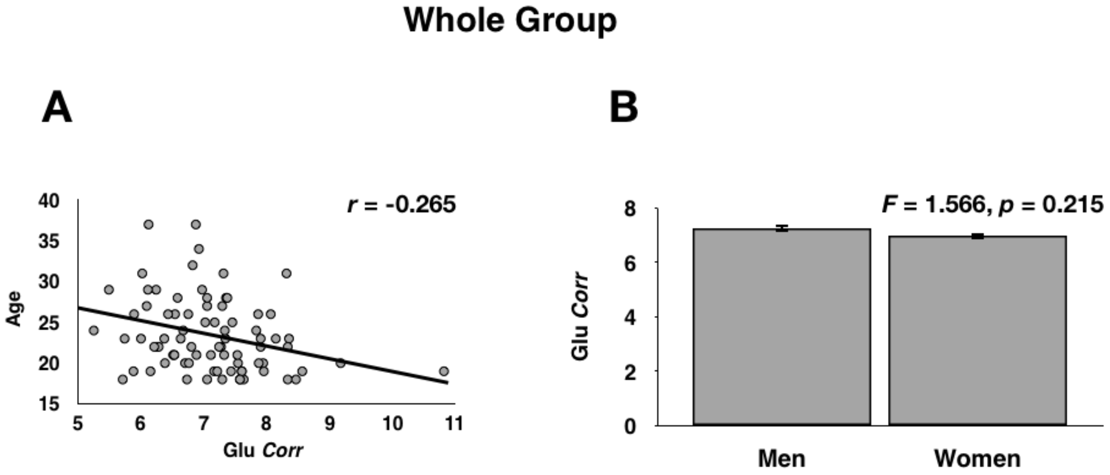


**Figure S2. A:** *Significant negative correlation between age and glutamate concentrations in all participants.* **B:** *Glutamate concentrations for both men and women; note that glutamate did not differ between genders.*

Glu *Corr* values did not correlate with the amount of cannabis smoked per day in the whole group ($r$ = -0.020, $p$ = 0.910, $BF$ = 0.192) (Figure S3), or separately in smokers ($r$ = 0.171, $p$ = 0.596, $BF$ = 0.501), intermittent smokers ($r$ = -0.157, $p$ = 0.534, $BF$ = 0.587) or non-smokers ($r$ = 0.228, $p$ = 0.591, $BF$ = 0.515).

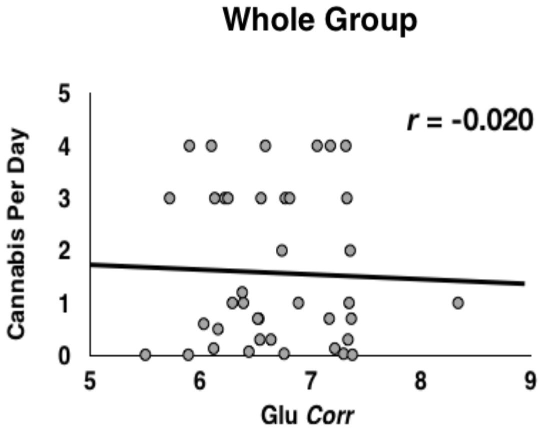


**Figure S3.** *Non-significant relationship between cannabis smoked per day and glutamate concentrations in all participants. Note that data points reflect values for only those participants who smoke more than zero cigarettes per day (i.e. daily and intermittent smokers only), for clarity.*

There was also a main effect of smoking status on Creatine Corr values (F(2,77) = 17.126, p < 0.001, $P_{BC}$ < 0.001, BF = 42.189), NAA Corr values (F(2,77) = 11.548, p < 0.001, $P_{BC}$ < 0.001, BF = 18.835) and MI Corr values (F(2,77) = 15.626, p < 0.001, $P_{BC}$ < 0.001, BF = 33.876) (see Fig. 2B). Subsequent analyses revealed that compared to non-smokers, daily smokers exhibited lower Creatine Corr values (F(1,54) = 13.745, p < 0.001, $P_{BC}$ = 0.002, BF = 24.784), MI Corr values (F(1,54) = 10.063, p = 0.002, $P_{BC}$ = 0.007, BF = 13.895) and NAA Corr values (F(1,54) = 11.034, p = 0.002, $P_{BC}$ = 0.005, BF = 15.249). Further, compared to non-smokers, intermittent smokers also exhibited lower Creatine Corr values (F(1,60) = 13.297, p < 0.001, $P_{BC}$ = 0.001, BF = 22.958), MI Corr values (F(1,60) = 13.146, p < 0.001, $P_{BC}$ = 0.001, BF = 20.690) and NAA Corr values (F(1,60) = 12.538, p= 0.001, $P_{BC}$ = 0.002, BF = 14.844). However, these values did not differ between daily and intermittent smokers (all ps > 0.507, all BFs < 0.328).

There was no effect of smoking status on all of the remaining 15 metabolites: alanine ($F$(2,77) = 0.221, $p$ = 0.728; $BF$ = 0.388), ascorbate ($F$(2,77) = 0.482, $p$ = 0.611; $BF$ = 0.537), aspartate ($F$(2,77) = 0.889, $p$ = 0.413; $BF$ = 0.612), GABA ($F$(2,77) = 0.875, $p$ = 0.421; $BF$ = 0.610), glucose ($F$(2,77) = 1.387, $p$ = 0.263; $BF$ = 1.039), glutamine ($F$(2,77) = 0.111, $p$ = 0.895; $BF$ = 0.212), glycine ($F$(2,77) = 0.925, $p$ = 0.398; $BF$ = 0.717), glutathione ($F$(2,77) = 0.436, $p$ = 0.648; $BF$ = 0.519), glycerophosphocholine ($F$(2,77) = 0.103, $p$ = 0.902; $BF$ =

0.196), phosphocholine ($F(2,77) = 0.448$, $p = 0.658$; $BF = 0.511$), lactate ($F(2,77) = 0.738$, $p = 0.509$; $BF = 0.603$), *N*-acetylaspartateglutamate ($F(2,77) = 1.452$, $p = 0.240$; $BF$ = 1.083), phosphorylethanolamine, ($F(2,77) = 0.692$, $p = 0.497$; $BF = 0.889$) scyllo-inositol ($F(2,77) = 0.184$, $p = 0.851$; $BF$ 229), taurine ($F(2,77) = 0.263$, $p = 0.708$; $BF = 0.362$).

In the whole group, glutamate concentrations were correlated with concentrations of creatine ($r = 0.790$, $p < 0.001$, $BF = 66.372$), *N*-acetylaspartate ($r = 0.852$, $p < 0.001$, $BF$ = 76.559), myo-inositol ($r = 0.723$, $p < 0.001$, $BF = 63.457$), GABA ($r = 0.500$, $p < 0.001$, $BF$ = 33.928), aspartate ($r = 0.577$, $p < 0.001$, $BF = 38.282$), and *N*-acetylaspartateglutamate ($r = 0.390$, $p < 0.001$, $BF = 19.369$), but not with concentrations of any of the remaining metabolites (all $p$s > 0.321, all $BF$s < 0.542).

Daily smokers, intermittent smokers and non-smokers did not differ in terms of Cramer-Rao lower bounds ($F(2,78) = 0.795$, $p = 0.455$, $BF = 0.385$), line width (in Hz) ($F(2,78)$ = 0.871, $p = 0.422$, $BF = 0.327$) or the signal-to-noise ratio ($F(2,78) = 0.167$, $p = 0.846$, $BF$ = 0.122). Finally, MRS data did also not differ between study 1 and study 2 in terms of Cramer-Rao lower bounds ($F(1,79) = 0.567$, $p = 0.570$, $BF = 0.264$), line width (in Hz) ($F(1,79) = 1.033$, $p = 0.313$, $BF = 0.398$) or the signal-to-noise ratio ($F(2,78) = 0.699$, $p$ = 0.406, $BF = 0.361$).
